# Antifungal exposure can enhance *Candida glabrata* pathogenesis

**DOI:** 10.1101/2025.07.31.667834

**Authors:** Gabriela Fior Ribeiro, Weronika Danecka, Logan Tomlinson, Edward W.J. Wallace, Delma S. Childers

## Abstract

Azole antifungal drugs directly inhibit lanosterol 14-ɑ-demethylase and indirectly affect the expression of metabolic, transmembrane transporter, and cell wall organization genes in fungal pathogens. It is not known how these indirect azole effects depend on dose, timing, and specific azole used, or how they influence host interactions. *Candida glabrata* (recently renamed *Nakaseomyces glabratus*) is the second leading cause of candidiasis, and clinical strains have high rates of intrinsic resistance to azoles. We investigated the early responses of reference strains BG2 and CBS138 to sub-inhibitory doses of fluconazole and voriconazole, and particularly, how these responses affect host-pathogen interactions. Cell wall profiling and transcriptomic data revealed highly similar responses for each strain to both azoles, including the upregulation of several virulence factors, such as yapsins. We also observed significant increases in CBS138 survival in macrophages and increased virulence in *Galleria mellonella* after voriconazole exposure. Using a combination of pharmacological inhibition of calcium ion channels and deletion strains, we determined that voriconazole-enhanced virulence requires a yapsin protease, *YPS1*, and is regulated via the calcineurin pathway and the cell wall integrity pathway, both of which regulate *YPS1* expression. We also observed that voriconazole treatment significantly reduced the virulence of the *bck1*Δ strain in *G. mellonella*, suggesting that inhibitors of the cell wall integrity pathway might potentiate azole activity by improving susceptibility to host killing. Our study provides new insight into short-term azole adaptation in *C. glabrata*, and importantly demonstrates that sub-inhibitory azole exposure can induce virulence factors and alter fungal pathogenesis.

**Article summary:** Antifungal drugs indirectly affect essential fungal cell processes, but we lack an understanding of how drug-induced changes affect fungal pathogenesis. We investigated how *Candida glabrata* adapts when exposed to azole drugs in terms of cell wall and transcriptional changes. Reference strains had similar transcriptional changes in response to azoles, but azole-treated CBS138 survived better in immune cells and caused more host death than untreated cells, suggesting that short-term azole treatment can significantly affect pathogenesis. Voriconazole-enhanced disease requires the calcineurin and cell wall integrity pathways and the virulence factor, *YPS1*, but could be blocked by a calcium ion channel inhibitor.

**Graphical Abstract:** 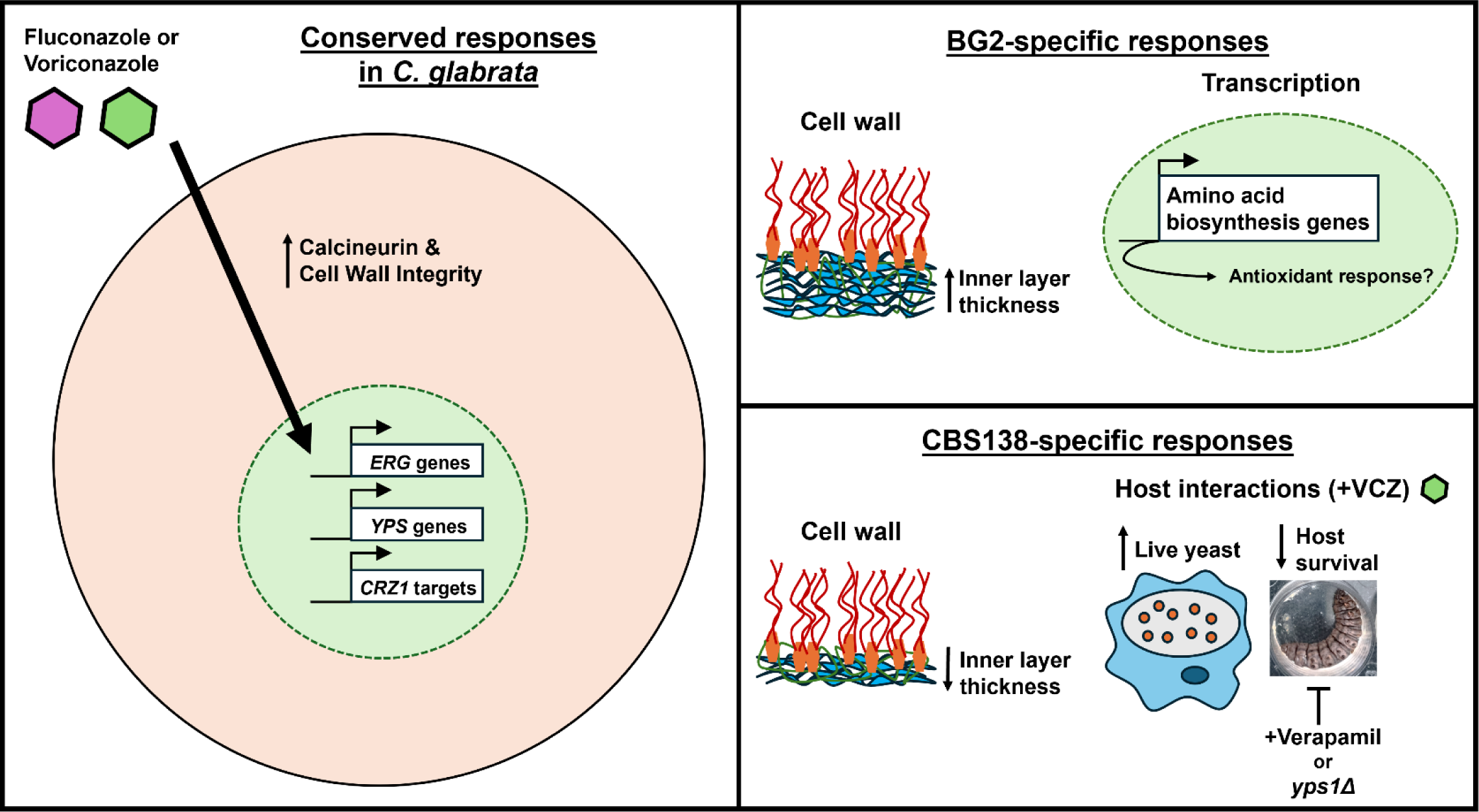

## Introduction

There is growing awareness and understanding of the importance of antifungal drug resistance and related phenomena in fungal pathogens, including tolerance, persistence and heteroresistance(Amich et al., 2025; Yang & Berman, 2024). These phenotypes have concerning implications for clinical disease management and treatment failure, which has driven investigations to understand the molecular mechanisms that permit pathogenic fungi to adapt to antifungals. Resistance is classically linked to stable genetic mutations that allow survival in high drug concentrations (Marie & White, 2009). However, other adaptive mechanisms, like tolerance and heteroresistance, have proven more difficult to characterize due to their transience within the population and lack of a clear, causal link to genetic modifications (Berman & Krysan, 2020; Rosenberg et al., 2018; Yang & Berman, 2024). Further, while antifungal resistance mutations are known to affect *Candida* species cell fitness and virulence (Bohner et al., 2022), we have a poor understanding of how other antifungal adaptive processes influence fitness and host-pathogen interactions.

*Candida glabrata* (recently renamed *Nakaseomyces glabratus*) is a major human fungal pathogen and the second leading cause of systemic candidiasis. *C. glabrata* is categorised as a high priority pathogen by the World Health Organisation (WHO, 2022) due to its serious clinical burden and high rates of antifungal resistance. Consistent with this classification, a recent Public Health England (PHE) surveillance study reported 17% and 21% of *C. glabrata* bloodstream clinical isolates as resistant to fluconazole and voriconazole, respectively (Budd et al., 2023). In comparison, only 1% of clinical isolates of the leading cause of candidiasis, *Candida albicans*, were resistant to fluconazole or voriconazole in the same study (Budd et al., 2023). Despite these high levels of azole resistance, fluconazole or voriconazole are sometimes still prescribed to patients with suspected fungaemias (Helmstetter et al., 2022).

Azoles directly inhibit lanosterol 14-ɑ-demethylase (encoded by *ERG11* in *Candida* species) leading to reduced ergosterol production, toxic sterol intermediate accumulation, and altered membrane fluidity. These processes also indirectly affect the expression of carbohydrate metabolism, transmembrane and ion transporters, and cell wall organization genes (Ribeiro et al., 2022). Little is known about the early cellular adaptations that pave the way for *C. glabrata* survival and drug resistance development in the host. We expect, based on existing transcriptomics and proteomics datasets, that the indirect effects of azole exposure on cell wall organisation and other biological processes alters host interactions. However, the effects of azole treatment on *C. glabrata* host-pathogen interactions are unclear.

In this study, we investigated the early adaptative responses of two *C. glabrata* reference strains, BG2 and CBS138, to sub-inhibitory doses of fluconazole and voriconazole, and particularly, how these responses affect host-pathogen interactions. We expected, based on our previous review of -omics datasets and the similarities in both drug class and reported resistance rates by PHE, that voriconazole and fluconazole might exert similar effects on cell wall remodeling and host-pathogen interactions (Budd et al., 2023; Ribeiro et al., 2022). While transcriptional and cell wall profiling data highlighted similar responses for each strain to both azoles, we unexpectedly observed that CBS138 survival in macrophages and virulence in *Galleria mellonella* infection studies was significantly improved after voriconazole exposure. Fluconazole pre-exposure also mildly enhanced both BG2 and CBS138 virulence in *G. mellonella*. We further demonstrated that voriconazole-enhanced virulence in CBS138 is at least partially dependent on the virulence factor and yapsin, *YPS1*, and the pathways required for *YPS1* expression, including calcium ion channel signaling, the calcineurin pathway, and the Slt2-MAPK (PKC) cell wall integrity pathway. Importantly, our study demonstrates how short-term adaptation to antifungals can induce survival strategies that enhance fungal pathogenesis.

## Materials and Methods

### Strains and Growth Conditions

*C. glabrata* reference strains BG2 and CBS138 (ATCC2001) (Cormack & Falkow, 1999; Dujon et al., 2004; Koszul et al., 2003; Schwarzmuller et al., 2014) were maintained by sub cultivation on YPD plates (2% glucose, 2% bactopeptone, 1% yeast extract) at 37°C from a frozen stock (-80°C).

Before each experiment, yeast cells were conditioned overnight in 5 or 25 mL MOPS-buffered liquid RPMI-1640 medium (final concentration: 2% glucose, MOPS 0.165 mol/L, pH 7) (Sigma R6504) at 37°C, 200 rpm. Yeast cells were back-diluted from overnight cultures (1×10^6^ cells/mL) for a further 4 hours growth in 10, 50 or 100 mL RPMI-1640 with antifungals (MIC_50_ concentrations, Table 1), 50 µg/mL verapamil (Sigma), or DMSO (Sigma) only according to experimental requirements.

**Table 1.** Minimum inhibitory concentrations for indicated antifungals and strains determined by broth microdilution method.

| Strain | Fluconazole |  | Voriconazole |  |
| --- | --- | --- | --- | --- |
|  | MIC<sub>50</sub> | MIC<sub>80</sub> | MIC<sub>50</sub> | MIC<sub>80</sub> |
| BG2 | 16mg/L | >64mg/L | 0.25mg/L | 4mg/L |
| CBS138 | 8mg/L | >16mg/L | 0.125mg/L | >2mg/L |

### Minimum Inhibitory Concentration (MIC) Determination

MIC testing was performed according to EUCAST guidelines ((The European Committee on Antimicrobial Susceptibility Testing, 2020). Briefly, *C. glabrat*a yeast cells were grown overnight in MOPS-buffered RPMI-1640 at 37°C. Cells were then centrifuged for 5 minutes, 7000 rpm, and the pellet was resuspended in RPMI-1640. 1×10^5^ yeast cells were added to each well in a 96 well plate in 90 μL RPMI-1640 and 10 μL of the respective drug dilution. Drug test ranges were 0.125-64 mg/L for fluconazole and 0.0156-8 mg/L for voriconazole. Plates were incubated for 24 hours at 37°C in the dark. The plates were then read on a spectrophotometer (VersaMax, SoftMax® Pro 7 Software), OD_530_, and the MIC_50_ and MIC_80_ for each strain and drug combinations were determined as the lowest concentration of drug needed to inhibit 50% or 80%, respectively, of cell growth.

### Flow Cytometry

To analyse cell wall carbohydrate exposure, cells grown with and without antifungals were inactivated overnight in 50 mM thimerosal (Sigma). Cells were then washed three times with PBS and counted by haemocytometer. 2.5×10^6^ cells were stained with 0.5 µg/mL Fc-Dectin-1 (kindly provided by Gordon Brown, MRC-CMM) and 1:200 diluted goat anti-human IgG antibody conjugated to Alexa Fluor 488 (Invitrogen), 50 µg/mL Wheat Germ Agglutinin (WGA) conjugated to Alexa Fluor 680 (Invitrogen), and 25 µg/mL Concanavalin A (ConA) conjugated to Texas Red (Invitrogen). Data were acquired for a minimum of 20,000 events on the Attune NxT (Thermo Fisher) and analysed using FlowJo v10 software (TreeStar Inc.) and gated as previously described (Ribeiro et al., 2025).

### Transmission Electron Microscopy (TEM)

Yeast cells were grown overnight in 5 mL RPMI-1640 (37°C, 200 rpm), counted, and 1×10^8^ cells were back-diluted and grown for a further 4 hours in 100 mL RPMI-1640 at 37°C, 200 rpm, containing voriconazole (MIC_50_) or DMSO (solvent control). Cells were then centrifuged at 4,000 rpm for 5 minutes. The concentrated pellet was placed between the sides of a small copper holder, enough to fill up the required space. Cells were then frozen in a high-pressure freezer and rapid transport system (Leica EMPACT2). Freeze substitution was carried out following the program detailed in Supplemental Table 2. Samples were then removed and placed in 10% Spurr’s (TAAB):Acetone for 72 hours – 30% Spurr’s overnight; 50% Spurr’s for 8 hours; 70% Spurr’s overnight; 90% Spurr’s for 8 hours. Subsequently, samples were embedded in Spurr’s resin at 60°C for at least 24 hours. Then 90nm sections were prepared using a diamond knife (Diatome Ltd, Switzerland) onto copper grids (EMResolutions) using a Leica UC6; and with Uranyless (TAAB) and Lead Citrate in a Leica AC20. Samples were viewed on the Transmission Electron Microscope JEM 1400 plus (JEOL) and captured using an AMT UltraVUE camera (AMT). Image J (Fiji) was used to measure the thickness of the inner (chitin and glucan) and outer (mannan) cell wall of 19-30 cells/group and 10-13 measurements/cell.

### BMDM Challenge

Bone Marrow-Derived Macrophages (BMDMs) were isolated from the femurs and tibias of male 12-weeks old C57BL/6 mice as previously described (Davies & Gordon, 2005; Gonçalves & Mosser, 2015). Mice were a kind gift from Gordon Brown and were randomly selected from in-house breeding colonies housed under specific-pathogen-free conditions at University of Aberdeen. Mice were not subjected to any regulated procedures prior to cervical dislocation and femur removal in accordance with ethical regulations approved by the University of Aberdeen Animal Welfare and Ethical Review body and the ARRIVE guidelines. BMDMs were maintained and differentiated in Dulbecco’s modified Eagle’s medium (DMEM; Sigma) supplemented with 10% heat inactivated Fetal Calf Serum (Gicbo), 15% L929 cell conditioned medium, 1% L-glutamine (Sigma), and 1% Penicillin/Streptomycin (Sigma). For macrophage interaction studies, 3×10^4^ BMDMs were plated on flat bottom 96-well plates and incubated overnight at 37°C, 5% CO2. BG2 and CBS138 wild-type cells were grown overnight in RPMI-1640, counted and 1×10^7^ cells were grown for a further 4 hours with MIC_50_ fluconazole, voriconazole, or <1% DMSO in 10 mL RPMI-1640 (2% Glucose, pH 7) at 37°C. Yeast cells were then washed with PBS and added to the 96-well plates in duplicate wells at a multiplicity of infection (MOI) of 3:1 (yeast cells to macrophages). Two hours post challenge the supernatant was removed, each well was washed with DMEM, and new media was added to remove unengulfed yeast cells. For the 2-hour timepoint 100 μL of 0.02% chilled SDS (Sodium Dodecyl Sulphate, Melford) was added to each well, its contents were scraped, serially diluted, spotted on YPD agar plates and incubated at 37°C. The same BMDM lysis and yeast recovery procedure was performed after 24 hours co-culture to determine CFU/mL and fold change in yeast cell recovery.

### *Galleria mellonella* Infection, Survival and Melanization

*G. mellonella* larvae were purchased from Livefood UK Ltd. (Axbridge, UK) and stored in wood shavings in the dark at room temperature prior to infection. *C. glabrata* yeast cells were grown overnight in 6 mL RPMI-1640 at 37°C, 200 rpm, and back-diluted to 1×10^8^ yeast cells in 100 mL RPMI-1640 with the indicated antifungals at MIC_50_ concentration or equivalent volume solvent, for a further 4 hours growth at 37°C, 200 rpm. Cells were washed and resuspended in sterile PBS. Larvae (∼250 mg weight) were randomly allocated into groups (specific sample sizes indicated on figure legends) and infected in the last left proleg with 5×10^6^ cells in a 50 μL/larvae suspension using a U-100 30G Micro-fine syringe (BD). Control groups were injected with 50 μL sterile saline only. Larvae were incubated at 37°C in the dark and survival and melanisation were assessed daily for a period of 6 days (144 hours). Larvae were scored for melanization as described previously (Usher et al., 2023). Briefly, larvae were considered partially melanised when their natural colour had been visibly altered, however they still did not present a fully darkened body. Larvae were considered fully melanised when their colour had been completely altered to a dark grey/brown pigmentation.

### Statistical Analyses

Statistical analyses were performed using GraphPad Prism v5.0 software (GraphPad Software) and IBM SPSS Statistics v27.0 (IBM Corp.). Specific experimental analyses described on figure legends. Macrophage-yeast survival and flow cytometry were analysed by Two-way ANOVA with Dunnett’s multiple comparisons test. TEM measurements were analysed by Two-Way ANOVA with Sidak’s multiple comparisons test. *G. mellonella* survival was analysed by Kaplan-Meier, Log-Rank pairwise over strata. A p value of <0.05 was considered to be significant, and the results are shown as mean ± standard error of the mean (SEM).

### RNA sequencing

Strains were streaked to single colonies on YPD agar plates for 2 days, then a single colony inoculated for each biological replicate into 5 mL RPMI-1640 with 2% glucose (Sigma R6504) and grown overnight. The next day, 10^8^ cells were pelleted and inoculated into 50 mL of RPMI-1640 media with 2% glucose pre-warmed to 37°C with drug or DMSO (mock) treatment. For drug addition, stock solutions of VCZ or fluconazole were prepared in DMSO, and the solution mixed with RPMI media immediately before inoculation. Antifungals were added at MIC_50_ concentration: VCZ at 0.25 ug/mL or 0.125 ug/mL for BG2 and CBS138, respectively; FCZ at 16 ug/mL or 8 ug/mL for BG2 and CBS138, respectively. Mock-treatment was performed using 0.004% DMSO, and DMSO was added to all VCZ and FCZ treatments to the final concentration of 0.004%.

After inoculation, cells were grown for 4 hours at 37°C with 200 rpm shaking, and harvested by pelleting cells for 3 min at 4000 rpm, the supernatant was decanted, and the cells were pelleted for 2 min at 3000 g to remove all supernatant, and the pellet frozen in liquid nitrogen, and stored at -80°C. Three biological replicates were prepared on successive days.

RNA was extracted using a modified silica column protocol following bead-beating with zirconia beads. The yeast pellets were thawed briefly on ice, then transferred to a screw cap tube and 200 μL of zirconia beads were added. 400 μL of RNA binding buffer (R1013, Zymo Research) was added, and the mixture was kept on ice for 1 minute. The tubes were transferred to PreCellys homogenizer (Bertin Technologies) and lysed using the following protocol: 10 seconds vortexing, followed by 10 seconds of waiting, repeated 3 times. The cells were then transferred to ice for 1 minute. Vortexing and incubation on ice were repeated a total of 6 times. The tubes were then centrifuged at 12,000 × g for 2 minutes. The supernatant was transferred to a Zymo Spin IIICG column (C1006, Zymo Research) and centrifuged at 12,000 g for 1 minute. 400 μL of ethanol was added to the flow through, mixed, transferred to a Zymo Spin IIC column (C1011, Zymo Research) and centrifuged for 1 minute. The column was washed with DNA/RNA Prep buffer (D7010-2, Zymo Research) and centrifuged for 1 minute, and then washed with DNA/RNA Wash buffer (D7010-3, Zymo Research) and centrifuged for 1 minute twice. The column was then transferred to a clean 1.5 mL tube, and RNA was eluted by adding 30 μL of water and centrifuging at 10,000 × g for 1 minute. The concentration of the samples was measured using Denovix. The quality and integrity of RNA was assessed on Fragment Analyzer (Agilent) using High Sensitivity RNA Kit (DNF-472-1000, Agilent). RQN and 28S/18S ratio were used to determine the quality of the sample.

RNA sequencing libraries were prepared from 500 ng total RNA using QuantSeq 3′ mRNA-Seq V2 Library Prep Kit REV (Lexogen, Vienna, Austria), a method that sequences a single fragment per mRNA, at the 3′ end proximal to the poly(A)-tail. Libraries were sequenced on NextSeq 2000 (Illumina, San Diego, USA)

RNA-Seq FASTQ files were processed using a Nextflow pipeline for QuantSeq data which is available online in GitHub (https://github.com/DimmestP/nextflow_paired_reads_pipeline). Software versions used were (Nextflow 3.1, FastQC 0.12.1, Cutadapt 4.3, HISAT2 2.2.1, SAMtools 1.17, MultiQC 1.14, BEDTools 2.30.0, Subread /FeatureCounts 3.11.3, DESeq2 1.40.2, Python 3.11.3, R 4.3.2, NCBI Datasets CLI 16.0.0) (Danecek et al., 2021; Di Tommaso et al., 2017; Ewels et al., 2016; Kim et al., 2019; Liao et al., 2014; Love et al., 2014; Martin, 2011; O’Leary et al., 2024; Quinlan & Hall, 2010; Wingett & Andrews, 2018). The reads were aligned to the genome sequence, for strain BG2: GCA_014217725.1; For CBS138, GCF_000002545.3. For assigning 3′ fragments to mRNAs, we used stranded alignment with an annotation file including 300nt added to the 3′ end of the annotated CDS. Differential gene expression was performed using DESeq2 (Love et al., 2014), R (R Core Team, 2021) and packages from the tidyverse (Wickham et al., 2019), and code is shared in GitHub (https://github.com/ewallace/cglab_rnaseq/). Genes were called as differentially expressed if they showed at least 2-fold difference at an adjusted p-value of 0.05 (5% false discovery rate), unless otherwise stated.

## Results

### Cell wall polysaccharide exposure is affected by sub-inhibitory azole treatment

We first investigated inhibitory concentrations of fluconazole (FCZ) and voriconazole (VCZ) for *C. glabrata* with the aim of identifying concentrations that impose significant stress while mimicking treatment failure (i.e. failure to completely inhibit or kill cells). We performed minimum inhibitory concentration (MIC) testing in accordance with EUCAST guidelines (The European Committee on Antimicrobial Susceptibility Testing, 2020). The concentration required to inhibit 50% of growth (MIC_50_) for CBS138 in FCZ and VCZ was 8 mg/L and 0.125 mg/L, respectively (Table 1). In comparison, the MIC_50_ for BG2 was 16 mg/L FCZ and 0.25 mg/L VCZ (Table 1), suggesting CBS138 is mildly more susceptible than BG2 to azole inhibition. This susceptibility was more pronounced for the MIC_80_ (concentration of drug required to inhibit at least 80% of growth compared to the control), where BG2 was four times more resistant to FCZ (64 mg/L versus 16 mg/L) and twice as resistant to VCZ (4 mg/L versus 2 mg/L) compared to CBS138 (Table 1).

Previous studies demonstrated that antifungal treatment leads to differential cell wall gene expression and significant changes in the cell wall that can have paradoxical effects on survival in *C. albicans* mammalian infections (Hopke et al., 2018; Lee et al., 2012; Walker et al., 2008). Therefore, we next tested whether short-term (4-hour) MIC_50_ antifungal exposure affected yeast cell wall polysaccharide detection among the *C. glabrata* reference strains, BG2 and CBS138. Cell wall features in BG2 were not majorly affected by FCZ or VCZ pre-treatment (Fig. 1a-c). In CBS138, there was a minor increase in β-glucan (∼1.21 and 1.27 fold-change for FCZ and VCZ, respectively) and chitin (∼1.23 and ∼1.16 fold-change for FCZ and VCZ, respectively) exposure levels in response to both azoles (Fig. 1a and 1c) compared to untreated cells. Mannan exposure was also significantly higher for CBS138 compared to BG2 (∼1.45 and ∼1.48 fold-change for FCZ and VCZ, respectively) (Fig. 1b).

**Figure 1.**
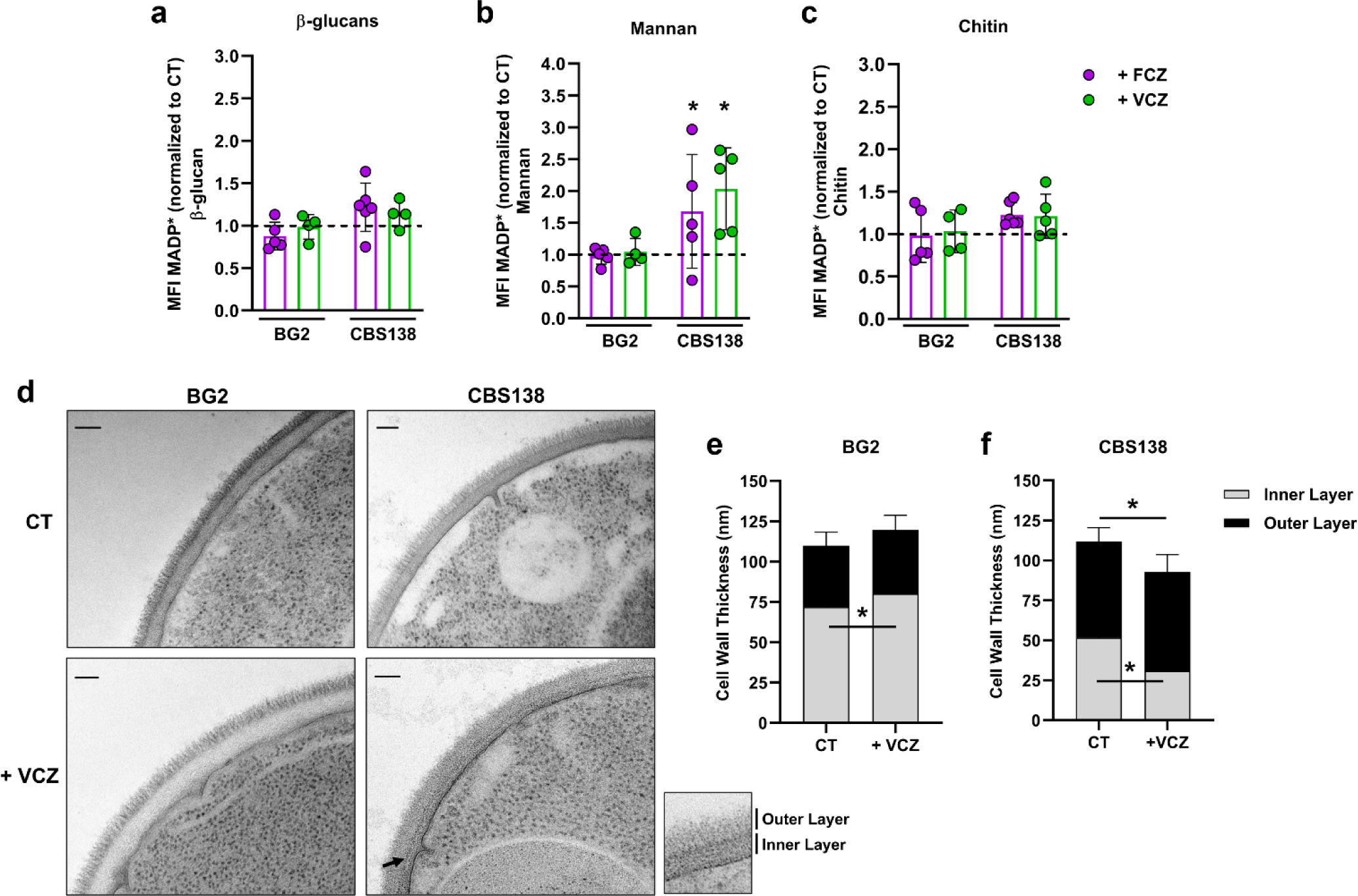
Pre-exposure to fluconazole and voriconazole impacts yeast cell wall architecture for *C. glabrata* reference strains BG2 and CBS138. We analyzed Median Fluorescence Intensities (MFI), based on Median Absolute Deviation of flow cytometry data, for *C. glabrata* reference strains BG2 and CBS138, pre-exposed or not to MIC_50_ fluconazole (FCZ) or voriconazole (VCZ), for β-glucan (Fc-Dectin-1) (a), mannan (Concanavalin A, ConA) (b) and chitin (Wheat Germ Agglutinin, WGA) exposure (c). Data represents two independent experiments, n = 4-6 biological replicates/group plotted as mean ± standard error of the mean (SEM) and normalized to their respective controls (DMSO only). * p ≤ 0.05 between indicated groups and their respective controls. Statistical analyses were done by Two-Way ANOVA with Dunnett’s multiple comparisons test. (d) Transmission Electron Microscopy (TEM) comparison of the cell wall of BG2 and CBS138. (e, f) TEM measurements of inner and outer cell wall thickness for BG2 (e) and CBS138 (f), pre-exposed or not to FCZ or VCZ. Scale bars represent 100 nm. Arrow indicates separation of inner and outer layers for CBS138 pre-treated with VCZ. n = 19-28 cells/group, 10-13 measurements/cell (maximum of 256 values plotted). Data plotted as mean ± SEM. * p ≤ 0.05 between indicated groups. Statistical analyses were done by Two-Way ANOVA with Sidak’s multiple comparisons test. CT, control (DMSO only).

Cell wall layer measurements by TEM further show that BG2 yeast cells pre-exposed to VCZ had slight differences in inner and outer wall thickness. BG2 cells had a slight, but significant, increase in inner (∼1.12 fold-change) layer thickness, but similar outer layer size (∼1.05 fold-change) compared to its control (Fig. 1d and 1e). However, and in contrast to BG2, CBS138 cells pre-treated with VCZ showed significantly reduced inner (∼0.6 fold change; p<0.05) but similar outer (∼1.03 fold-change) layer thickness compared to controls (Fig. 1d and 1f).

Taken together, our flow cytometry data showed minimal changes in β-glucan and chitin exposure after azole treatment which was unexpected given the changes in inner layer thickness by TEM. However, both azoles increased mannan exposure in CBS138 compared to BG2 (∼1.9-fold; Fig. 1b), and our TEM measurement data indicates that CBS138 cells generally had a thicker outer cell wall layer compared to BG2 (Fig. 1d-f).

### Voriconazole exposure enhances CBS138 pathogenesis

We observed above that short-term FCZ and VCZ exposure differentially impacted some carbohydrate exposure and the gross cell wall architecture of the two *C. glabrata* reference strains, CBS138 and BG2. Cell wall composition plays an important role in modulating host responses, including fungal clearance by immune cells (Gow et al., 2017). Therefore, we next tested how azole pre-exposure affected yeast survival in macrophages by measuring yeast colony forming unit (CFU) recovery following macrophage challenge (Ribeiro et al., 2025).

As before, yeast cells were treated with or without MIC_50_ FCZ or VCZ prior to co-incubation with bone marrow-derived macrophages. After 2 hours of yeast-macrophage challenge, we observed no statistically significant differences in yeast recovery from macrophages between azole-treated and untreated groups for either strain, though there was a trend toward a greater percentage recovery of the CBS138 inoculum from groups pre-exposed to azoles, especially VCZ (Fig. 2a and 2b). After 24 hours of co-incubation, we still observed no significant differences in yeast recovery between treatment groups for BG2 (Fig. 2c). However, at 24 hours we recovered significantly higher CFUs of VCZ-treated CBS138 cells compared to FCZ-treated cells or the control, suggesting that VCZ-treated CBS138 cells were able to replicate better within macrophages than FCZ-treated and untreated yeast cells (Fig. 2c). As expected, based on the CFUs recovered at each time point, the fold change in yeast recovery between 2 and 24 hours showed no variance for BG2 between groups and a trend of increased survival for VCZ-treated CBS138 yeast cells compared to FCZ-treated and untreated cells (Figure 2d).

**Figure 2.**
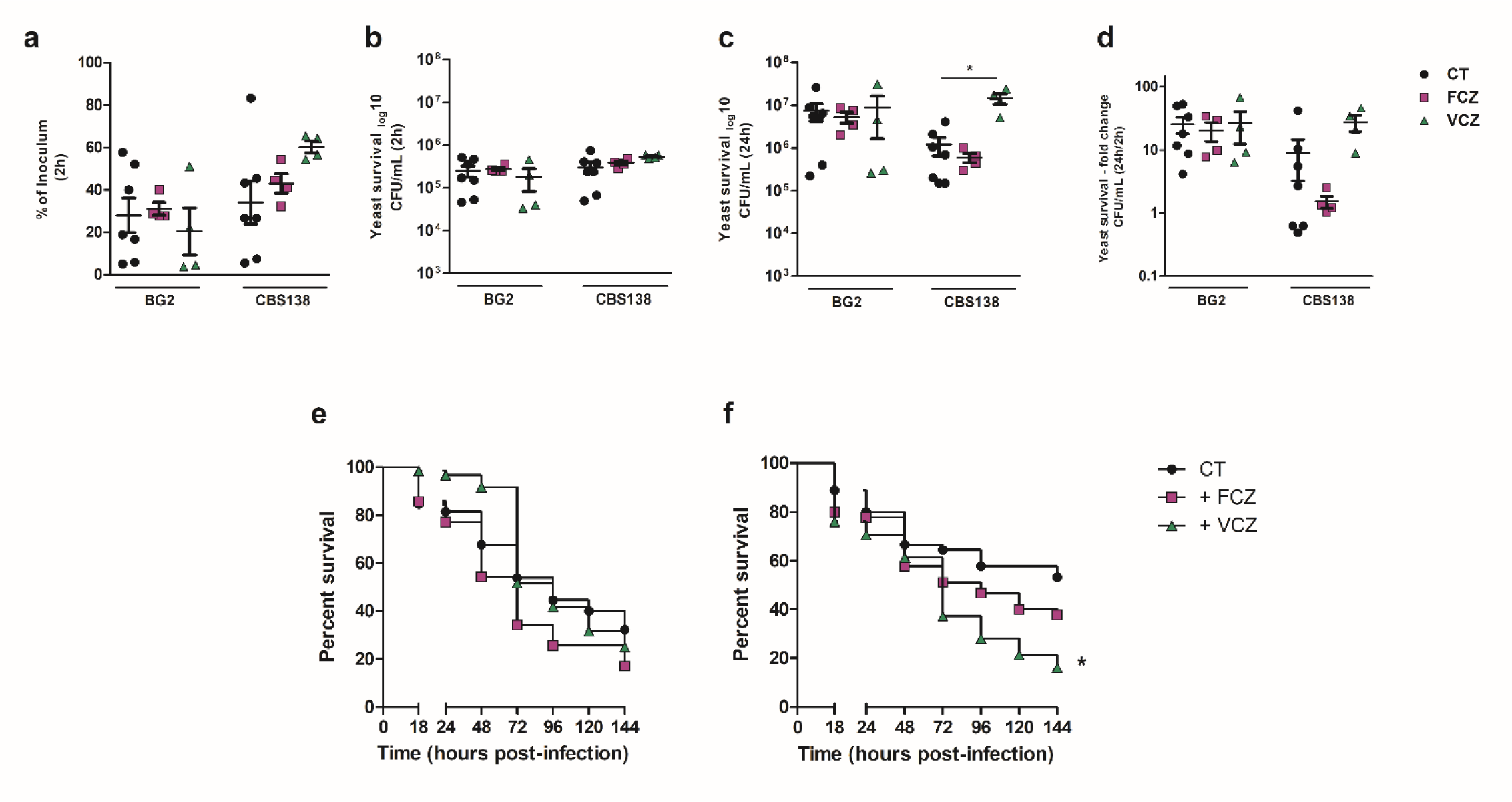
VCZ pre-exposure improves CBS138 yeast cell recovery after 24-hour challenge with BMDMs and enhances virulence in *G. mellonella*. (a-d) BMDMs were challenged in technical duplicate at an MOI of 3:1 *C. glabrata* cells to macrophages. Internalized yeast cells at 2 hours post-challenge are presented as percent of initial inoculum (a) and CFU/mL (b). (c) Internalized yeast cells were also determined at 24 hours post-challenge. (d) The fold change of yeast survival was determined by the ratio of recovered cells at 24 hours vs 2 hours post-challenge. The mean and SEM are indicated by the line and whiskers on each plot. Data represents six independent experiments, n = 4-8 biological replicates per group for macrophage yeast survival, mean of technical replicates. Statistical analyses were done by Two-Way ANOVA with Dunnett’s multiple comparisons test. * p ≤ 0.05 between indicated groups. (e and f) *G. mellonella* larvae were injected with 5×10^6^ BG2 (e) or CBS138 (f) yeast cells that had been exposed to no treatment, FCZ or VCZ for 4 hours. Survival was monitored for up to 144 hours post-infection. Data represents five independent experiments n = 10-15 larvae per group per experiment. * p ≤ 0.05 between the indicated group versus control. Statistical analyses were done by Kaplan-Meier. CT, DMSO only control.

We next tested whether azole pre-treatment affected *C. glabrata* virulence in the *G. mellonella* systemic infection model. Consistent with our macrophage interaction data, we observed no significant differences in *G. mellonella* survival during infection with azole-treated and untreated BG2 yeast cells (Fig. 2e). For CBS138, infection with FCZ-treated yeast induced slightly faster larval death than the control, but VCZ-treated yeast killed larvae significantly faster than untreated cells (Fig. 2f; p<0.05) with a final survival difference >40% between control and VCZ-exposed infection groups.

Altogether, our findings suggest that azole pre-treatment has minimal effects on BG2 host interactions, but azoles, and especially VCZ, trigger enhanced survival and virulence in CBS138.

### Transcriptomic responses to azole drugs are broadly similar across strains

We designed an RNA-seq experiment to identify transcriptomic changes that might explain differences in azole-enhanced virulence between BG2 and CBS138. As above, we treated yeast for 4 hours with either FCZ or VCZ at MIC_50_ concentration or a mock-treated DMSO-only control. We prepared 3 biological replicates and made libraries using a 3′ mRNA-Seq approach. Extracted RNA was high-quality and the aligned reads passed all relevant quality checks, including high correlations between replicate samples (Figs S1, S2).

Transcriptome profiles clustered both by strain and by drug treatment, as revealed by principal component analysis of the regularized log-counts (Fig 3a). Comparing principal components 1 and 2 shows that differences between strain and drug are almost orthogonal (Fig 3a) and contain almost 70% of the variance (Fig 3b). Transcriptome profiles from treatment by VCZ and FCZ were very similar within each strain, both in the principal component plot (Fig 3a) and by correlation analysis (Figs S1, S2). Differential gene expression analysis confirmed these findings: hundreds of genes were significantly differentially expressed between strains in each growth condition, and differentially expressed between drug-treated cells and mock-treated cells for each strain (Fig S3). However, within-strain no significantly differentially expressed genes were found between FCZ and VCZ (Fig S3). We conclude that as each strain received the same “subjective dose” of VCZ and FCZ, at the strain-specific MIC_50_ concentration, the transcriptomic responses to these two azoles were practically indistinguishable.

**Figure 3.**
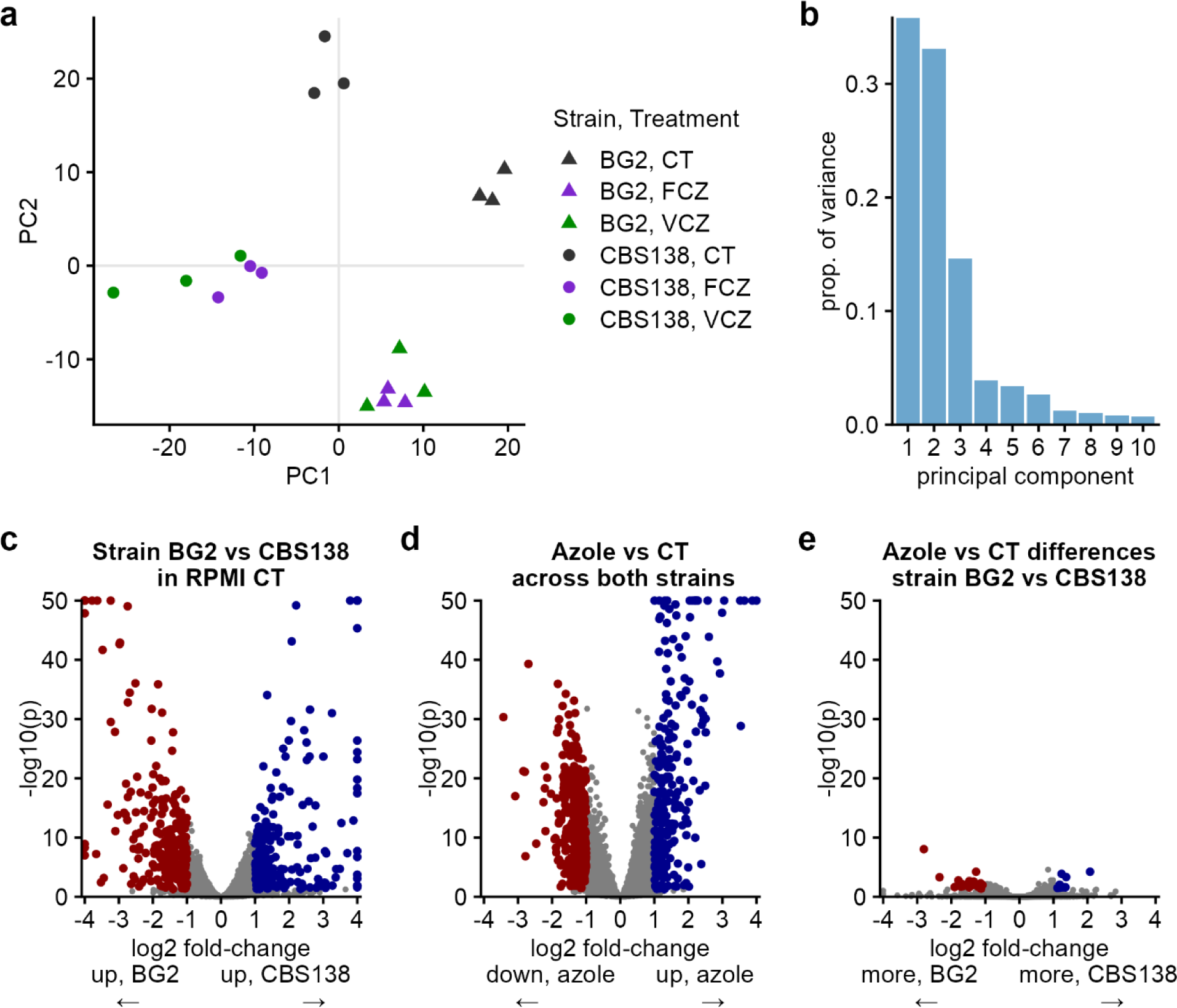
Azole drugs induce a consistent transcriptomic response, overlaid on strain-dependent baseline gene expression. (a) Principal Component Analysis shows that RNA-seq samples cluster by strain and by azole drug treatment. Principal components 1 and 2 of the regularized logarithm of counts per gene are shown for all samples; see methods for details. (b) Between-sample variance is concentrated in principal components 1 and 2, that panel a shows cluster by strain and azole treatment. (c) There are extensive baseline gene expression differences between strains BG2 and CBS138. (d) A consistent azole-dependent transcriptomic response is identified by pooling azole-dependent differential expression across both FCZ and VCZ drugs across both strains BG2 and CBS138. (e) There are minimal strain-dependent differences in drug-induced gene expression, detected using the interaction term in a DESeq2 analysis with both factors (‘design = ∼ Strain * Drug’). See methods for details of differential gene expression analysis across 3 biological replicates using DESeq2, and supplementary figure S3 for additional pairwise differential expression plots.

Thus, we break down the analysis into three main components: baseline differences between strains (Fig. 3c), common drug-regulated transcripts in both strains (Fig. 3d), and transcripts that were differentially induced in one strain compared to the other, i.e. drug-strain interactions (Fig. 3e).

The baseline differences between BG2 and CBS138 are extensive (Fig 3c). In the control samples, 194 genes had significantly higher expression in CBS138 compared to 265 in BG2 (Fig. 3c). Genes with higher expression in CBS138 are enriched in GO categories including those associated with cell wall assembly, cell-cell adhesion, and cell aggregation (i.e. *GAS3*, *SWM1*, *FKS3*, *EPA6*, *EPA3*, *ZAP1*, *KSS1*, and several uncharacterized genes), and some involved in amino acid biosynthetic processes including lysine and other amino acid biosynthesis (i.e. *LYS9*, *ARG1*, *IDP1*, *MET13*, *LYS12*, *LEU2*, *LYS21*, *STR3*, and several uncharacterized genes). Genes with higher expression in BG2 are enriched in a variety of categories related to metabolism including trehalose metabolism (*TPS2*, *ATH1*, *UGP1*, *CAGL0H02387g*, *CAGL0K03421g*), stress responses (including *GCN4*, *MSN4*, *YHB1*, *TUP11*, *NUC1*, *KRE29*, *TDH3*, *HSP12*, *SSA3*, *HSP78*), and translation (including *FRS2*, *TIF1*, *EFT2*). We did not find a clear picture here about how baseline transcriptomic differences between strains could explain their different phenotypes, so focused on azole responses subsequently.

Common azole-regulated targets are extensive (Fig 3d) and consistent with previous datasets (Ribeiro et al., 2022). The 258 significantly azole-induced upregulated genes are enriched in GO terms such as lipid metabolism, organelle organization, response to chemical, and vesicle-mediated transport. Consistent with azoles targeting ergosterol production, ergosterol biosynthesis pathway genes were induced in both strains including *ERG1*, *ERG2*, *ERG3*, *ERG5*, *ERG7*, *ERG11*, *ERG24*, and *ERG25* (Fig 4). Azole upregulates the multidrug resistance transcription factor *PDR1*, along with target transporters involved in drug resistance, *CDR1* and *PDH1*, but not the homolog *SNQ2* (Fig 4a). Several yapsins, proteases which are important virulence factors that suppress host immune responses (Rasheed et al., 2018), were upregulated in both strain backgrounds (Fig 4). The 355 significantly azole-downregulated genes include genes involved in ribosomal biogenesis and translation, consistent with drug treatment having a negative impact on growth. Full GO results are included in the online supplementary data.

**Figure 4.**
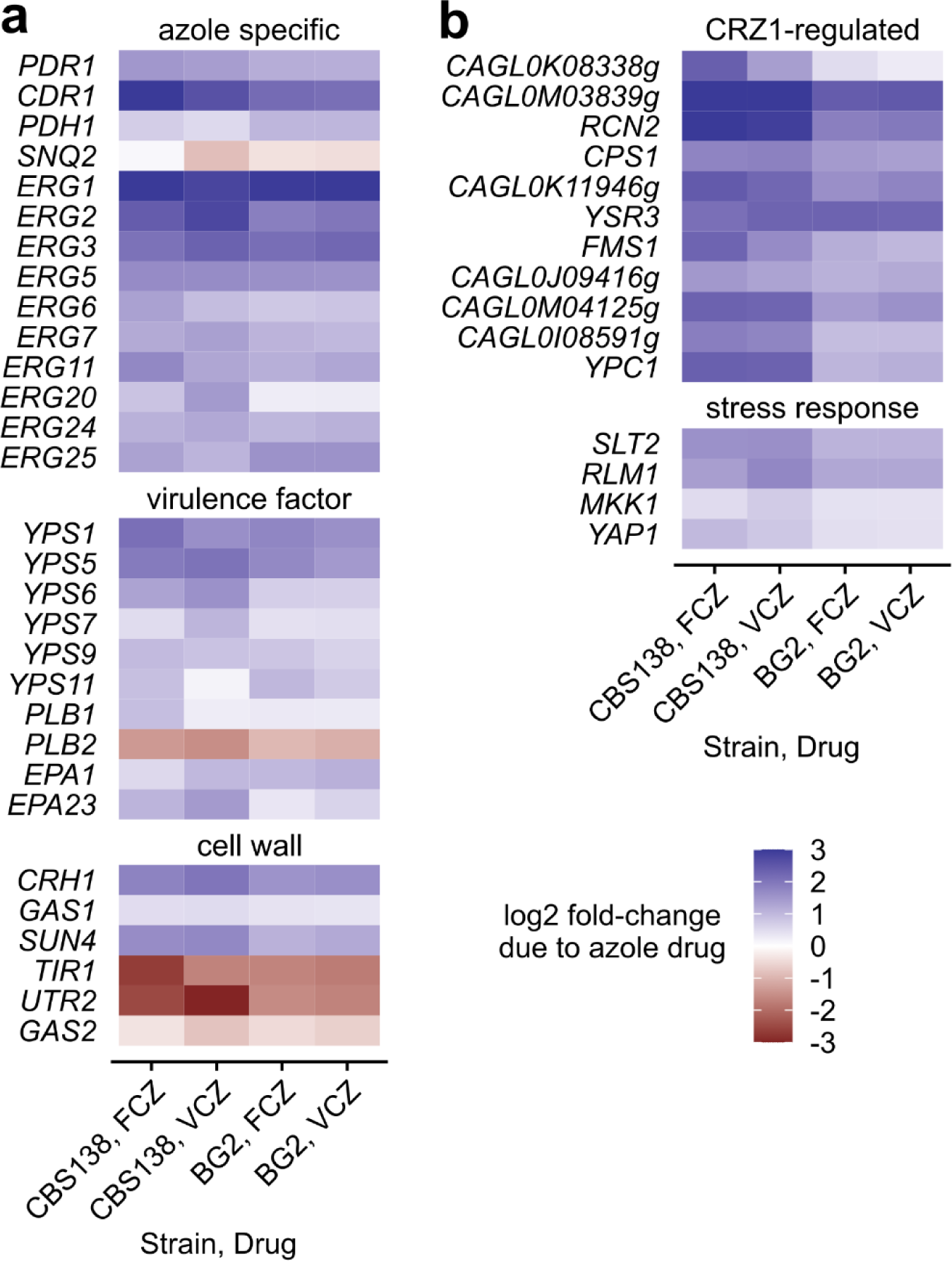
Azole drugs induce differential expression of specific genetic pathways. (a) Azoles induce expression of select multidrug transporters (*PDR1* transcription factor, *CDR1* transporter, *PDH1* transporter, but not the *SNQ2* transporter), along with multiple ergosterol biosynthesis genes. Azoles also induce expression of the yapsin family of aspartyl proteases. Azoles further induce differential expression of different cell wall genes. (b) Azoles induce expression of multiple genes in the *CRZ1* calcineurin-responsive transcription factor pathway, and other key stress response genes.

We expected that transcripts that are differentially induced by azoles in CBS138 compared to BG2 might explain the increase in virulence in azole-treated CBS138. Surprisingly, very few genes fall into this category: only 9 are more induced in CBS138 than BG2, and 22 vice versa, at a false discovery rate of 5% and minimal 2-fold expression change (Fig 3e). The genes induced more in CBS138 include the *YAP6* transcription factor, that has roles in stress responses (Merhej et al., 2016), and eight uncharacterized genes. The 22 genes induced more in BG2 are largely associated with transport and metabolic processes (full list available in online supplemental).

The calcineurin pathway and its transcription factor, *CRZ1*, are important regulators of azole resistance in *C. glabrata* (Vu et al., 2023) and provide a critical stress response to combat azole-mediated membrane disruption in *C. albicans* (Onyewu et al., 2004). In our datasets, we observed differential expression of *CRZ1*-dependent genes, including induction of the calcineurin negative feedback regulator, *RCN2*, in response to azole treatment. Given the diverse roles calcineurin plays in cell wall maintenance, stress responses and host interaction, we hypothesized that the enhanced virulence of azole-treated CBS138 requires calcineurin activity and induction of *CRZ1*-dependent targets, like yapsins.

### Calcium ion channel inhibition suppresses voriconazole-enhanced virulence

The calcineurin pathway is typically known for its role in calcium signaling, and blocking calcium channels alongside azole treatment synergistically inhibits the growth of drug-resistant *C. albicans* strains (Liu et al., 2016). We therefore tested the importance of calcium for azole-enhanced virulence using the drug verapamil to inhibit calcium-importing ion channels (Fig. 5a) (Teng et al., 2008; Yu, Q. et al., 2013).

**Figure 5.**
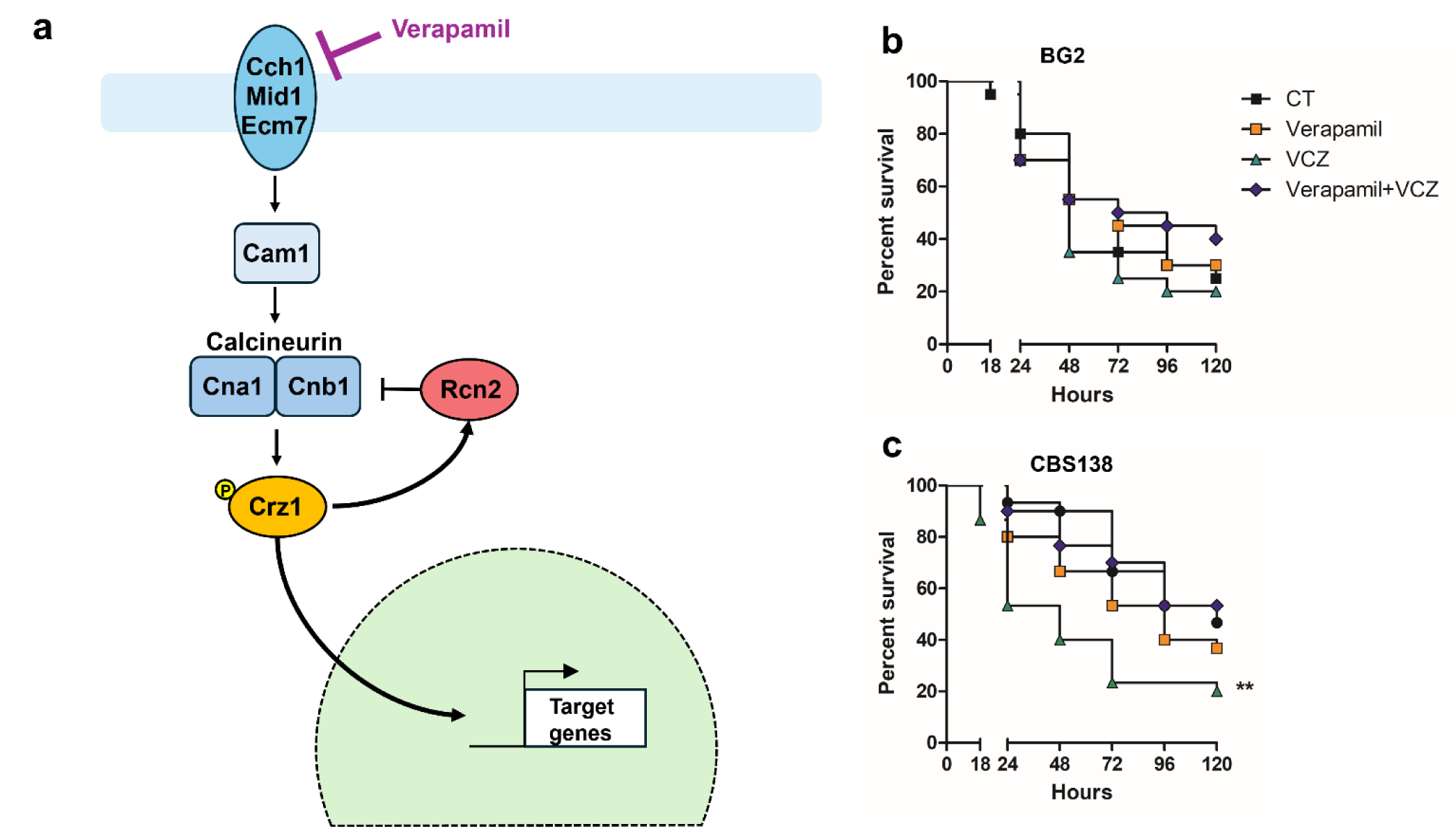
Verapamil inhibits voriconazole-enhanced virulence. (a) Diagram of the calcineurin pathway. (B-C) *G. mellonella* larvae were injected with 5×10^6^ BG2 (b) or CBS138 HTL (c) yeast cells that had been pre-exposed to no treatment, MIC_50_ VCZ, 50 µg/mL verapamil, or both VCZ and verapamil for 4 hours. Survival was monitored for up to 120 hours post-infection. Data represents three independent experiments n = 10 larvae per group per experiment. ** p ≤ 0.01 between the VCZ group versus control and VCZ+Verapamil. Statistical analyses were done by Kaplan-Meier. CT, DMSO only control.

CBS138 and BG2 cells were untreated or treated for 4 hours with 50 µg/mL verapamil, MIC_50_ VCZ, or a combination of both verapamil and VCZ prior to infecting *G. mellonella* (Fig. 5). Similar to our earlier observations (Fig. 2e), there were no significant differences in virulence for larvae infected with either VCZ-treated or untreated BG2 cells (Fig. 5b). Verapamil treatment alone also did not significantly alter BG2 virulence compared to untreated yeast. BG2 cells pre-treated with the combination of verapamil and VCZ resulted in slightly better overall larval survival (∼40% survival versus 25% for the control), but this was not significant compared to infection with untreated cells.

We again observed significantly enhanced virulence for CBS138 cells pre-treated with VCZ compared to untreated controls (p<0.01, Fig. 5c) Treatment with verapamil alone did not significantly alter CBS138 virulence (Fig. 5c). However, the combination of verapamil and VCZ rescued larval survival back to control levels (Fig. 5c), suggesting that calcium ion channels are required for voriconazole-enhanced virulence. Indirectly, azoles may be inadvertently triggering calcium signaling and other pathways in a way which promotes CBS138 virulence.

### Cell wall integrity, calcineurin and *YPS1* are necessary for voriconazole-enhanced virulence in CBS138

Our verapamil study suggested that target genes downstream of the calcineurin pathway may be important for voriconazole-enhanced virulence in CBS138. One target of this pathway that was upregulated in our RNA-Seq dataset is the yapsin, *YPS1*, which requires both calcineurin and cell wall integrity pathway (Slt2-MAPK) signaling for its expression (Fig. 6a) (Miyazaki et al., 2011). We used available mutants in a published gene deletion collection (Schwarzmuller et al., 2014) to test our hypothesis that calcineurin, the cell wall integrity pathway, and virulence factor *YPS1* contribute to voriconazole-enhanced virulence.

**Figure 6.**
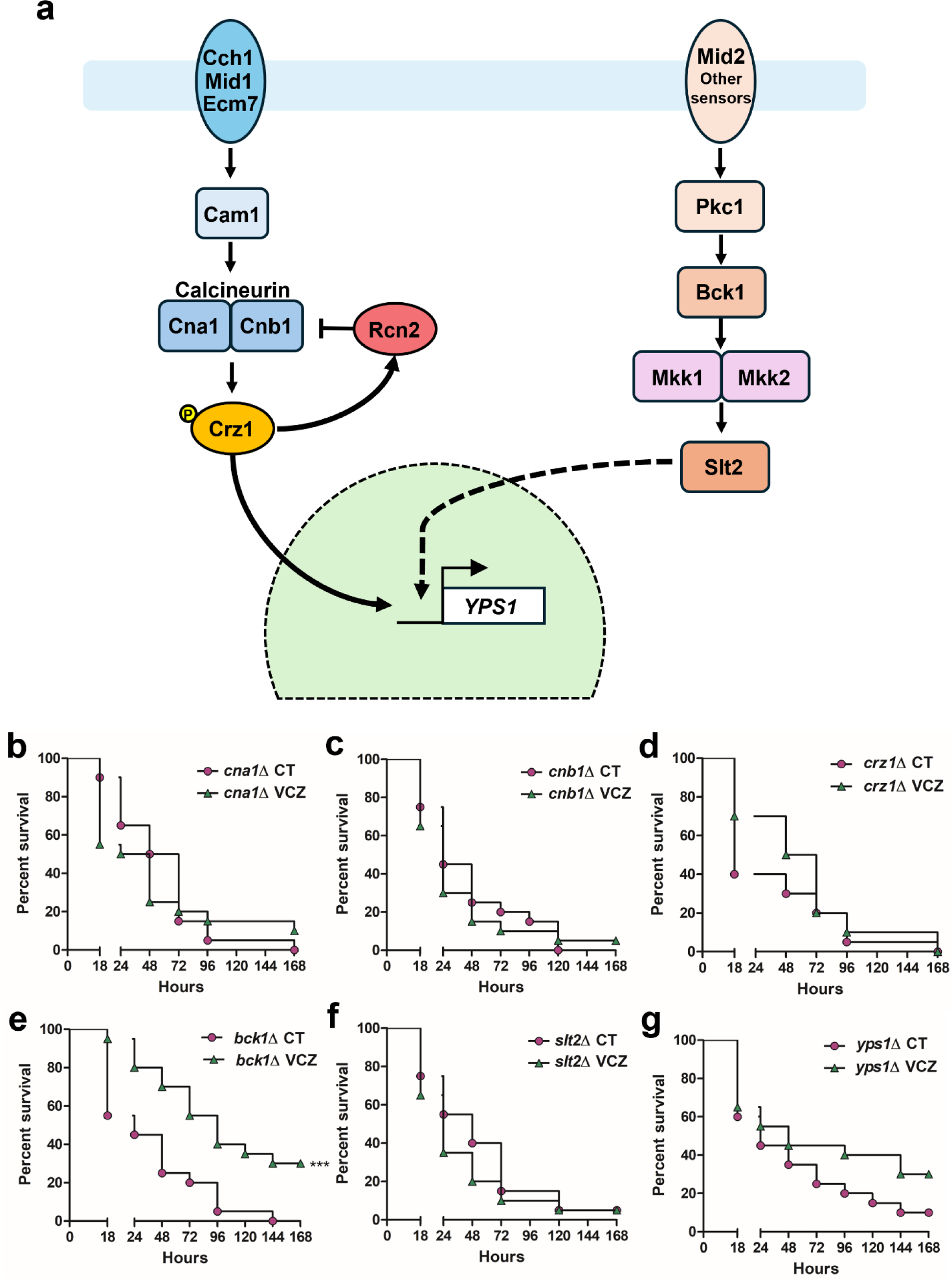
*YPS1*, calcineurin pathway, and PKC pathway components are required for voriconazole-enhanced virulence. (a) Diagram of calcineurin and MAPK pathways coordinating *YPS1* expression in *C. glabrata*. (b-g) *G. mellonella* larvae were injected with 5×10^6^ cells of the indicated strain that were untreated or pre-treated with MIC_50_ VCZ for 4 hours. Survival was monitored for up to 168 hours post-infection. Data represents two independent experiments n = 10 larvae per group per experiment. * p ≤ 0.05 between the VCZ group versus control. Statistical analyses were done by Kaplan-Meier. CT, DMSO only control.

*cna1*Δ, *cnb1*Δ, *crz1*Δ, *bck1*Δ and *slt2*Δ were grown for 4 hours with or without VCZ at MIC_50_ concentration prior to infecting *G. mellonella* (Fig. 6b-f). Larvae infected with either treatment group for *cna1*Δ, *cnb1*Δ, *crz1*Δ and *slt2*Δ showed no significant differences in survival at 168 hours post-infection. The only slight, but still statistically insignificant difference was for *crz1*Δ during early infection (18 hours), where we observed reduced survival for larvae infected with untreated cells compared to VCZ-treated cells (∼40% survival versus ∼70% survival, respectively; Fig. 6d). Surprisingly, VCZ-treated *bck1*Δ cells were attenuated for virulence compared to untreated cells. Untreated *bck1*Δ cells killed all larvae within 144 hours while infection with VCZ-treated cells resulted in ∼40% survival (Fig. 6f). Overall, these data support our hypothesis that Slt2-MAPK and the calcineurin pathway are necessary for the CBS138 voriconazole-enhanced virulence phenotype.

We next tested *yps1*Δ, which we grew with and without VCZ for 4 hours prior to infecting *G. mellonella* (Fig. 6g). Larvae infected with VCZ-treated or untreated *yps1*Δ cells died at similar rates up to 48 hours post-infection. After 48 hours, larvae infected with VCZ-treated *yps1*Δ cells died at a slower rate than larvae infected with untreated cells in a trend similar to the *bck1*Δ strain.

Altogether, our data suggest that the voriconazole-enhanced virulence we observed for CBS138 requires both the cell wall integrity and calcineurin signaling pathways and their downstream co-regulated virulence factor, *YPS1*.

## Discussion

Azole treatment stalls fungal growth by directly inhibiting ergosterol biosynthesis, but also indirectly affects cellular processes such as cell wall biogenesis (Ribeiro et al., 2022). We have a poor understanding of how the direct and indirect effects of azole treatment contribute to fungal fitness and persistence in the host. Most studies investigate antifungal responses using a single reference strain or drug. This approach is perfectly reasonable given potential issues with study feasibility and cost, but makes systematic comparisons between drugs, strains and species difficult. In our study, we address some of these issues by exploring the early adaptation of two *C. glabrata* reference strains to two azole drugs.

Published transcriptomics and proteomics datasets indicated that cell wall biogenesis processes are differentially regulated by multiple antifungal drug classes, including azoles (Ribeiro et al., 2022). The cell wall is the first point of contact between fungi and host cells, and cell wall carbohydrates are an important mediator of host innate immune responses. However, we observed few changes in cell wall carbohydrate exposure after 4 hours of FCZ or VCZ treatment (Fig 1a-c), though we did observe differences in inner cell wall thickness in VCZ-treated cells compared to controls and in CBS138 mannan exposure in response to both drugs (Fig 1d-f). Our flow cytometry probes include lectins specific for the inner cell wall carbohydrates chitin and β-1,3-glucan (Allen et al., 1973; Palma et al., 2006), therefore it is possible that changes in the inner cell wall architecture are related to β-1,6-glucan abundance, which we are currently unable to detect. The differences in cell wall thickness and CBS138 mannan exposure also suggest potential changes in cell wall protein abundance, though there are limited data available characterizing the fungal proteome in response to antifungal treatment (Pais et al., 2016; Ribeiro et al., 2022). Pais et al. found that clotrimazole treatment altered the abundance of 37 *C. glabrata* membrane proteins; however, 25 of these proteins had decreased abundance. In *C. albicans*, ketoconazole treatment increased the abundance of 32 proteins, but the protein isolation procedure was not specific to the cell wall (Hoehamer et al., 2010). Our RNA-Seq analysis shows evidence for differential regulation of cell wall-associated genes (Fig. 4a), but no clear bias towards mechanisms that would be consistent with our flow cytometry profiling, such as general upregulation of genes encoding mannoproteins.

Azole exposure and associated minor cell wall changes did not appear to affect host interactions for strain BG2. Azole-treated BG2 cells had no major differences in inoculum uptake by macrophages, intracellular survival, or virulence in *G. mellonella* versus untreated controls. However, we observed that sub-inhibitory azole treatment altered CBS138-host interactions. FCZ-treated cells had a mild trend towards decreased intracellular replication over 24 h and slightly increased virulence in *G. mellonella* compared to control cells. VCZ-treated cells had higher intracellular inoculum recovery at 2 h post-macrophage co-incubation, were recovered at significantly higher CFU after 24 h co-incubation with macrophages and were significantly more virulent in *G. mellonella* than control cells. While we do not yet know the reason behind these drug-specific effects in CBS138, others have shown that azoles can localize to multiple subcellular compartments, including the mitochondria (Benhamou et al., 2017; Elias et al., 2019; Koren et al., 2024), and may have off-target effects on heme production or other processes which could explain the differences in virulence phenotypes between drugs.

We performed RNA-Seq to determine what transcriptional changes might drive the strain- and drug-dependent differences we observed in *C. glabrata* virulence. We were surprised to discover that both azoles induced nearly indistinguishable transcriptional differences for BG2 and CBS138, with only 31 genes showing altered differential expression between strains in response to azoles. Most of these 31 genes are uncharacterized, and several genes upregulated in BG2 during azole exposure are associated with metabolic processes such as sterol uptake (*TIR3*), the tricarboxylic acid cycle (CAGL0L02079g, *ACO2*, CAGL0K11616g), and amino acid biosynthesis for methionine (*MET15*), lysine (CAGL0J06402g, CAGL0K07788g, *LYS9*, *LYS12*, *LYS21*) and arginine (*ARG8*, CAGL0I08987g). Notably, the azole-induced lysine biosynthetic genes are homologous to *S. cerevisiae* enzymes that play a vital role in cellular tolerance against oxidative stress (O’Doherty et al., 2014; Olin-Sandoval et al., 2019). Their upregulation in BG2 suggests that cells mount an oxidative stress defense early during azole treatment, which is consistent with studies demonstrating that FCZ and other azoles induce reactive oxygen species (ROS) in multiple *Candida* species (Gonzalez-Jimenez et al., 2023; Mahl et al., 2015). While azole-induction of the lysine biosynthetic pathway appeared to be BG2-specific, it is important to note that 4 out of 5 of these genes already had significantly elevated baseline expression in CBS138 under control conditions compared to BG2 (online supplemental).

Our RNA-Seq data also indicated that azoles induced the expression of several virulence factors, particularly the yapsins, in both reference strains. Yapsins are glycosylphosphatidylinositol (GPI)-linked aspartyl proteases that participate in cell wall remodeling, immune evasion, and virulence. *YPS1* and *YPS5* expression is regulated by *CRZ1* via the calcineurin pathway (Chen et al., 2012), and *YPS1* expression requires additional input from the cell wall integrity (Slt2-MAPK) signaling pathway (Miyazaki et al., 2011). The calcineurin and cell wall integrity pathways play important roles in drug and stress tolerance across fungal pathogens. In particular, the calcineurin pathway is required for azole and caspofungin tolerance (Yu, S. et al., 2015), thermotolerance (Chen et al., 2012), and maximizes *C. glabrata* survival in response to micafungin and manogepix treatment (Pavesic et al., 2024). We observed that several *C. glabrata CRZ1*-regulated targets were induced by both FCZ and VCZ, including the calcineurin negative feedback regulator, *RCN2* (Chen et al., 2012). The loss of VCZ-enhanced virulence for genetic mutants in the calcineurin pathway as well as after pharmacological inhibition of calcium ion channels using verapamil (Scorzoni et al., 2020; Teng et al., 2008) strongly suggest that the calcineurin pathway is an essential component of the CBS138 VCZ-enhanced virulence phenotype. Further, genes in the cell wall integrity pathway and *YPS1* were also necessary for CBS138 VCZ-enhanced virulence in *G. mellonella*. Unexpectedly, VCZ-treated *bck1*Δ was significantly less virulent than control cells, and VCZ-treated *yps1Δ* also had reduced virulence compared to the control, though this was not statistically significant. This virulence attenuation is not likely due to differences in drug cidality because we obtained similar CFU compared to the controls when inocula were plated to verify cell counts (data not shown). However, this virulence defect was not observed with *slt2Δ.* The VCZ-induced virulence defect for *bck1Δ* and *yps1Δ* suggests that *BCK1* and *YPS1* contribute to compensatory processes that are necessary for *C. glabrata* azole adaptation independently of *SLT2*. Further, previous work has shown that *YPS1* expression was not fully repressed in the *slt2*Δ background in response to thermal stress (Miyazaki et al., 2011), suggesting that low levels of *YPS1* expression are sufficient to maintain virulence post-VCZ treatment in the *slt2*Δ background.

In conclusion, our study provides fundamental insights into the baseline and azole-induced differences of two key *C. glabrata* reference strains. We demonstrated how azole-exposure alters CBS138 host interactions, which can be blocked by calcium ion channel inhibition (i.e. verapamil) or deletion of key components in the calcineurin and cell wall integrity pathways. Our study establishes that these pathways, in addition to contributing to cell survival and drug resistance development, also contribute to maintaining virulence in the host after azole treatment and offer the prospect that interfering with cell wall integrity signaling could potentiate azole-induced fitness defects in the host.

## Author contributions

G.F.R - experimental design, data acquisition and analysis, writing & editing; W.D. - experimental design, data acquisition and analysis, funding acquisition, writing & editing; L.T. - data acquisition and analysis, editing; E.W.J.W. - experimental design, data analysis, funding acquisition, writing & editing; D.S.C. - experimental design, data acquisition and analysis, funding acquisition, writing & editing

## Data availability

RNA-seq data are available on Gene Expression Omnibus (GEO), accession number GSE273379. Supplemental code and GO tables are available at: https://github.com/ewallace/cglab_rnaseq/.

## Acknowledgments

We are grateful to our colleagues at the University of Aberdeen Institute of Medical Sciences core facilities and wish to acknowledge Andrea Holme and the Iain Fraser Cytometry Centre and Debbie Wilkinson, Gillian Milne and Lucy Wight in the Microscopy and Histology Facility for training and assistance with cytometry and microscopy. We thank our Aberdeen Fungal Group colleagues, especially Donna MacCallum, for helpful discussions. We thank Wallace lab members for helpful discussions. We thank Richard Clarke, Angie Fawkes, and Lee Murphy for performing RNA-seq at the Genetics Core of the Edinburgh Wellcome Trust Clinical Research Facility. We also thank Jane Usher at University of Exeter for helpful comments and discussion.

