## Supplemental Data for "Antifungal exposure can enhance *Candida glabrata* pathogenesis"

**Supplementary figures**

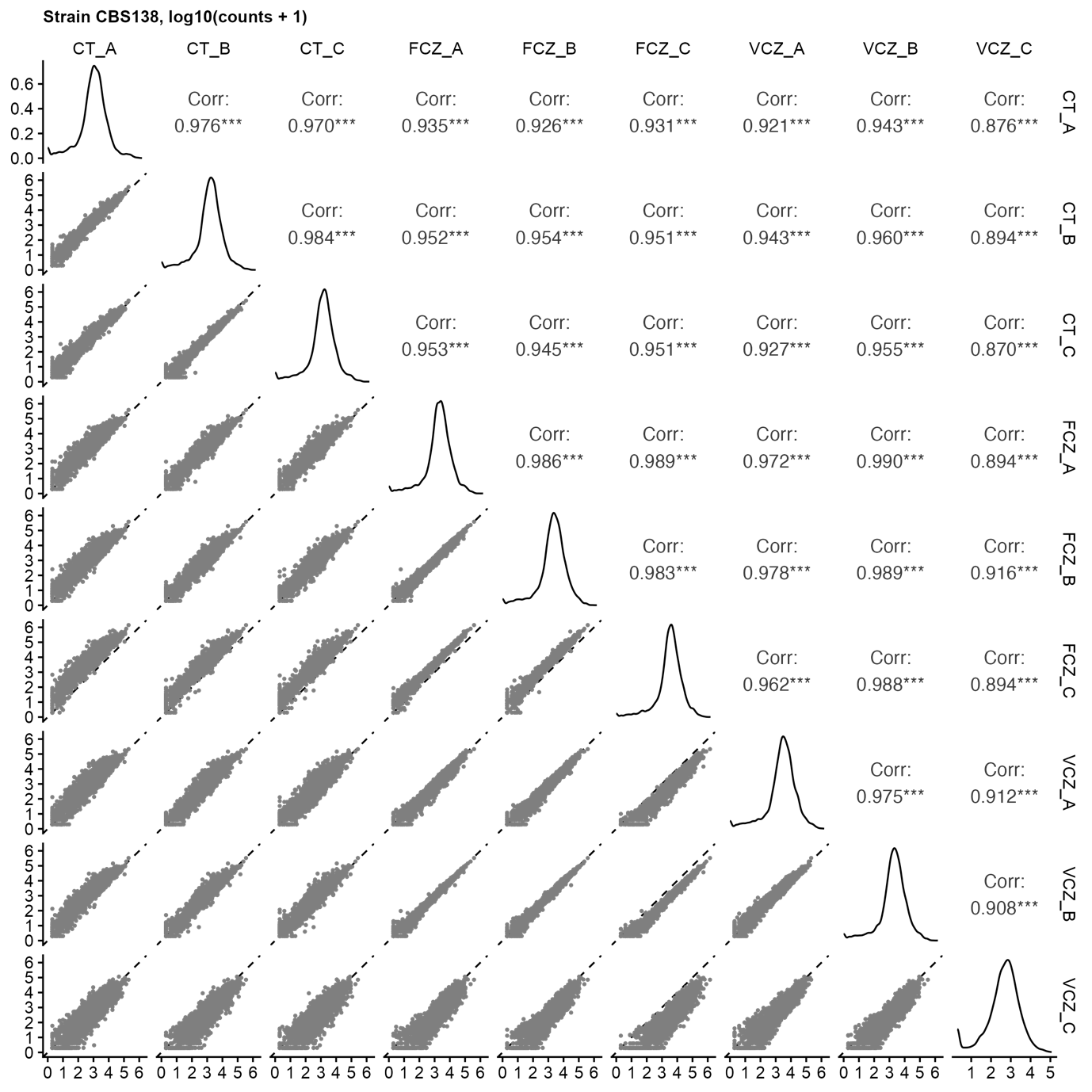

Fig S1. RNA-sequencing counts are highly reproducible. Figures show log10 (count per gene + 1) for all samples, and correlation coefficients, for strain CBS138.

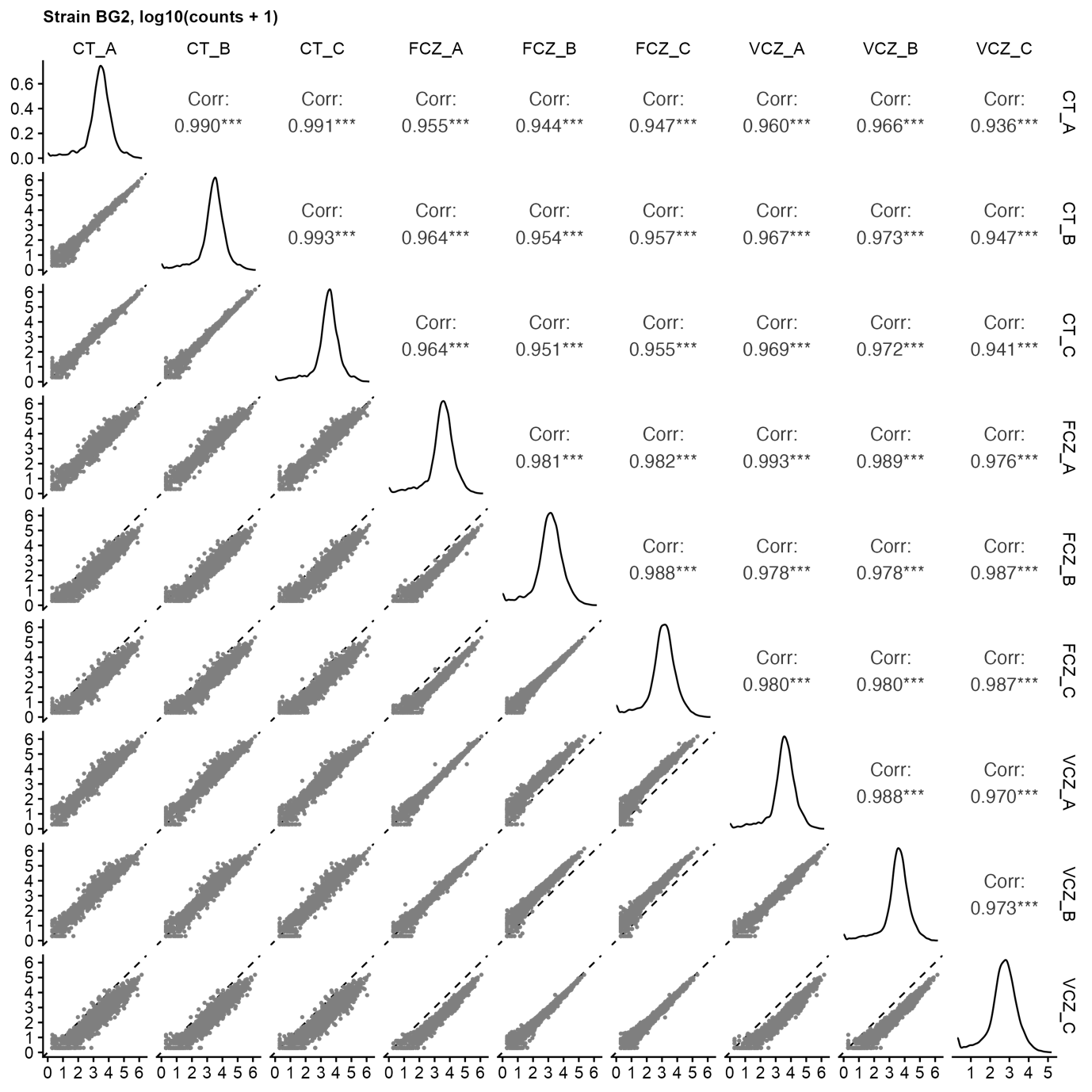

Fig S2. RNA-sequencing counts are highly reproducible, as Fig S1 but for strain BG2.

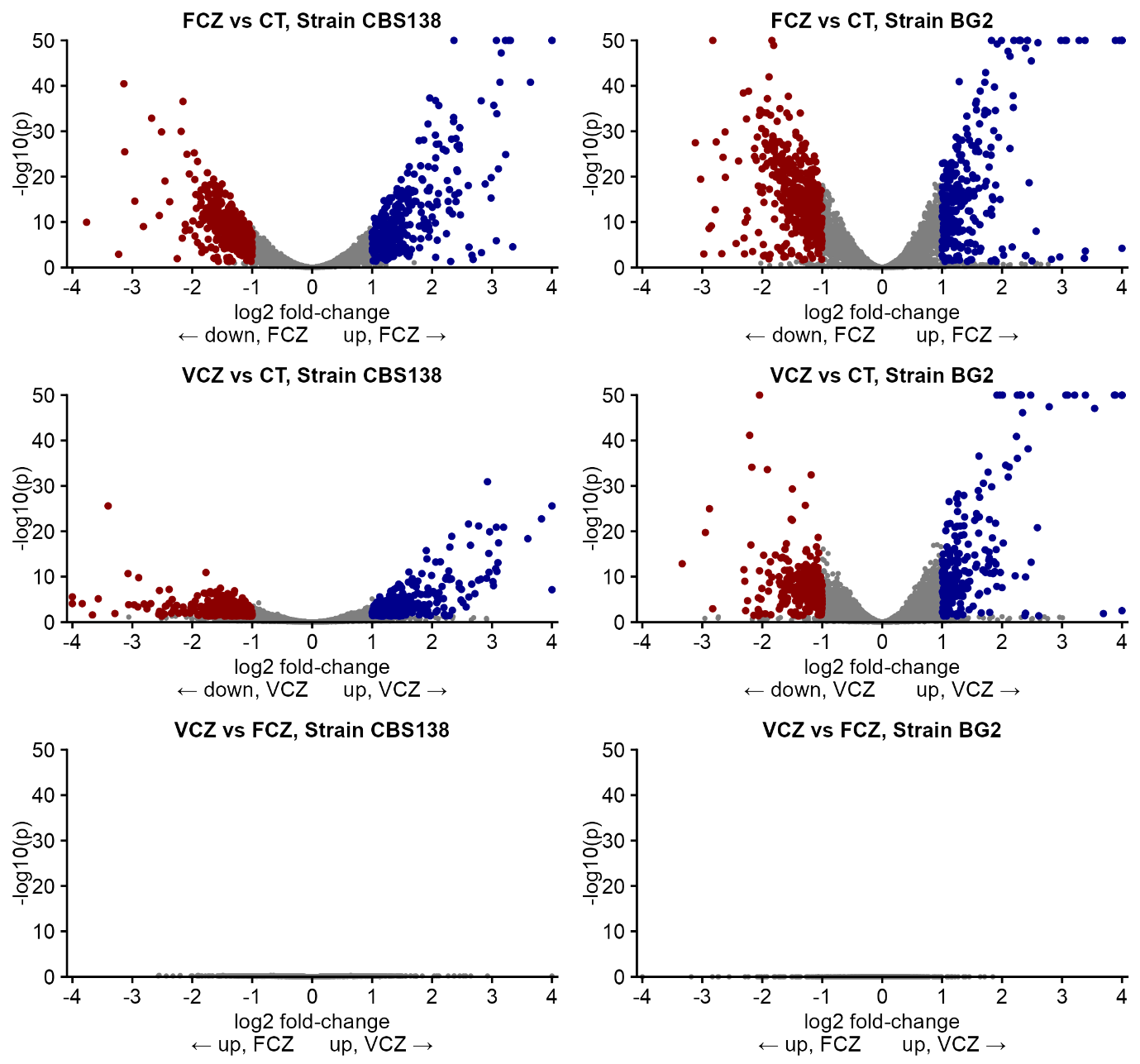

Fig. S3. Pairwise comparisons between drug treatments within-strain. Both FCZ and VCZ at MIC50 concentrations lead to extensive differential gene expression. However, there is no significant difference between FCZ- and VCZ- treated cells at MIC50-matched drug concentrations.

**Table S1. List of all strains used in the study.**

| **Strain** | **Genotype** | **Parent** | **Source** |
| --- | --- | --- | --- |
| **BG2** | *N. glabratus* Wild-Type | - | (Cormack & Falkow, 1999) |
| **CBS138 (ATCC2001)** | *N. glabratus* Wild-Type | - | (Dujon et al., 2004; Koszul et al., 2003) |
| **CBS138 HTL** | *his3*∆::FRT *leu2*∆::FRT *trp1*∆::FRT | CBS138 | (Schwarzmüller et al., 2014) |
| ***cna1*Δ** | *his3*∆::FRT *leu2*∆::FRT *trp1*∆::FRT *cna1Δ*::*NAT1* | HTL | (Schwarzmüller et al., 2014) |
| ***cnb1*Δ** | *his3*∆::FRT *leu2*∆::FRT *trp1*∆::FRT *cnb1Δ*::*NAT1* | HTL | (Schwarzmüller et al., 2014) |
| ***crz1*Δ** | *his3*∆::FRT *leu2*∆::FRT *trp1*∆::FRT *crz1Δ*::*NAT1* | HTL | (Schwarzmüller et al., 2014) |
| ***bck1*Δ** | *his3*∆::FRT *leu2*∆::FRT *trp1*∆::FRT *bck1Δ*::*NAT1* | HTL | (Schwarzmüller et al., 2014) |
| ***slt2*Δ** | *his3*∆::FRT *leu2*∆::FRT *trp1*∆::FRT *slt2Δ*::*NAT1* | HTL | (Schwarzmüller et al., 2014) |
| ***yps1*Δ** | *his3*∆::FRT *leu2*∆::FRT *trp1*∆::FRT *yps1Δ*::*NAT1* | HTL | (Schwarzmüller et al., 2014) |
